# Differential effects of novel kappa opioid receptor antagonists on dopamine neurons using acute brain slice electrophysiology

**DOI:** 10.1101/2020.04.24.059352

**Authors:** Elyssa B. Margolis, Tanya L. Wallace, Lori Jean Van Orden, William J. Martin

## Abstract

Activation of the kappa opioid receptor (KOR) contributes to the aversive properties of stress, and modulates key neuronal circuits underlying many neurobehavioral disorders. KOR agonists directly inhibit ventral tegmental area (VTA) dopaminergic neurons, contributing to aversive responses [1,2]; therefore, selective KOR antagonists represent a novel therapeutic approach to restore circuit function. We used whole cell electrophysiology in acute rat midbrain slices to evaluate pharmacological properties of four novel KOR antagonists: BTRX-335140, BTRX-395750, PF-04455242, and JNJ-67953964. Each compound concentration-dependently reduced the outward current induced by the KOR selective agonist U-69,593. BTRX-335140 and BTRX-395750 fully blocked U-69,593 currents (IC_50_ = 1.3 ± 0.9 and 4.6 ± 0.9 nM, respectively). JNJ-67953964 showed an IC_50_ of 0.3 ± 1.3 nM. PF-04455242 (IC_50_ = 19.6 ± 16 nM) exhibited partial antagonist activity (∼60% maximal blockade). In 50% of neurons, 1 μM PF-04455242 generated an outward current independent of KOR activation. BTRX-335140 (10 nM) did not affect responses to saturating concentrations of the mu opioid receptor (MOR) agonist DAMGO or the delta opioid receptor (DOR) agonist DPDPE, while JNJ-67953964 (10 nM) partially blocked DAMGO responses and had no effect on DPDPE responses. Importantly, BTRX-335140 (10 nM) rapidly washed out with complete recovery of U-69,593 responses within 10 min. Collectively, we show electrophysiological evidence of key differences amongst KOR antagonists that could impact their therapeutic potential and have not been observed using recombinant systems. The results of this study demonstrate the value of characterizing compounds in native neuronal tissue and within disorder-relevant circuits implicated in neurobehavioral disorders.

## Introduction

One of the major challenges in drug development is predicting whole animal responses based on pharmacological characterization in heterologous systems. Recent biological reports indicate that the effect of drugs on G protein coupled receptor function *in situ* in brain tissue is not reliably predicted from results in expression systems [3–8]. Therefore pharmacological characterizations made in brain tissue likely relate better to behavioral outcomes than those made in cell-based expression assays.

Interest in the kappa opioid receptor (KOR) as a target for therapeutic development has been growing consistently as clinical and preclinical studies have identified its role in aversive behavioral states. KOR agonists produce profound adverse effects in humans, specifically fatigue, sedation, confusion, impaired concentration, and anxiety. Furthermore at higher concentrations visual and auditory hallucinations and feelings of depersonalization have been reported [9,10]. Homologous effects have been described in animal models (reviewed in [11]). Finally, blockade or genetic deletion of the KOR significantly reduces aversive responses to stress [12–14], drug withdrawal [15–17], and pain [18], and has antidepressant-like effects [19] in preclinical models, suggesting that KOR selective antagonists could be useful therapeutic agents.

Historically, the known synthetic KOR antagonists, including the most widely used KOR antagonist for laboratory research norbinaltorphimine (norBNI), have properties limiting their clinical potential, including long lasting blockade of KOR agonist activity [20,21]. These long lasting effects have been alternatively attributed to prolonged retention time in the brain [22] or a signaling process involving the activation of the c-Jun N-terminal kinase (JNK) pathway [23,24]. In addition, some possess poor selectivity for KOR over other opioid receptors and have other off-target effects [25,26]. Recently, new compounds have been synthesized to overcome these limitations [27]. In particular, BTRX-335140 (1-(6-ethyl-8-fluoro-4-methyl-3-(3-methyl-1,2,4- oxadiazol-5-yl)quinolin-2-yl)-*N*-(tetrahydro-2H-pyran-4-yl)piperidin-4 amine) has been reported to have a medication-like duration of action [28] and is currently in clinical trials. PF-04455242 (2- methyl-N-((2’-(pyrrolidin-1-ylsulfonyl)biphenyl-4-yl)methyl)propan-1-amine; Pfizer) [29] and JNJ- 67953964 (formerly LY2456302 / CERC-501) (S)-3-fluoro-4-(4-((2-(3,5-dimethylphenyl)pyrrolidin-1-yl)methyl)phenoxy)benzamide) also have been or are in clinical development, respectively, as selective KOR antagonists.

In rats, the model species used here, KOR activation in the VTA is aversive and directly inhibits the activity of dopamine neurons [1,30]. Several brain regions that innervate the VTA express mRNA for the endogenous KOR ligand, dynorphin, including ventral striatum, amygdala, and lateral hypothalamus [31–33]. One hypothesis is that a major contributor to maladaptive aversiveness is dynorphin release from one or more of these inputs, inhibiting dopamine neuron activity that KOR antagonist treatment would reverse, generating relief [34–36]. Therefore, here we investigated properties of 4 synthetic KOR antagonists in VTA dopamine neurons, since KOR actions on the midbrain dopaminergic system have been implicated in some of the proposed clinical indications for KOR antagonist treatments [11]. We used an acute midbrain slice and whole cell electrophysiology preparation to evaluate the potency, selectivity, and reversibility of a selection of recently developed KOR antagonists to gain a better understanding of their pharmacologies in brain tissue.

## Materials and Methods

All animal protocols were conducted in strict accordance with the recommendations of the National Institutes Health (NIH) in the Guide for the Care and Use of Laboratory Animals. Research protocols were approved by the Institutional Animal Care and Use Committee (University of California at San Francisco, CA), approval ID AN169369-3B. Decapitation was performed after deeply anesthetizing the rats with isoflurane to minimize discomfort.

### Tracer injections

Male Sprague Dawley rats (27–29 d) were anesthetized with isoflurane and secured in a skull stereotax. A glass pipette tip attached to a Nanoject II (Drummond Scientific, Inc.) was stereotaxically placed in the medial prefrontal cortex (mPFC) (from bregma (in mm): anteroposterior (AP), +2.6; mediolateral (ML), ±0.8; ventral (V), − 4.0 from skull surface). Neuro- DiI (7% in ethanol; Biotium) was slowly injected, 200 nL per side. All injection sites were histologically examined and only data collected from rats with on target injections were included in the analysis.

### Slice preparation and electrophysiology

Most recordings were made in control male Sprague Dawley rats (28d–adult). Recordings in retrogradely labeled neurons were made 6–8 d after surgery. Horizontal brain slices (150 μm thick) were prepared using a vibratome (Leica Instruments). Slices were prepared in ice-cold aCSF (in mM: 119 NaCl, 2.5 KCl, 1.3 MgSO_4_, 1.0 NaH_2_PO_4_, 2.5 CaCl_2_, 26.2 NaHCO_3_, and 11 glucose saturated with 95% O_2_ – 5% CO_2_) and allowed to recover at 33°C for at least 1 h. Slices were visualized under a Zeiss AxioExaminer.D1 with differential interference contrast, Dodt, and near infrared optics, and epifluorescent illumination to visualize DiI-labeled neurons. Whole cell recordings were made at 33°C using 3–5 MΩ pipettes containing (in mM): 123 K-gluconate, 10 HEPES, 0.2 EGTA, 8 NaCl, 2 MgATP, 0.3 Na_3_GTP, and 0.1% biocytin (pH 7.2, osmolarity adjusted to 275). Liquid junction potentials were not corrected during recordings.

Recordings were made using an Axopatch 1-D (Molecular Devices), filtered at 5 kHz and collected at 20 kHz using custom procedures written for NIDAQ and IGOR Pro (National Instruments and WaveMetrics, respectively). In control animals, VTA neurons were selected from throughout the VTA. All experiments were completed in voltage clamp, V_holding_ = −60 mV. Input resistance and series resistance were measured once every 10 s with a 4 mV hyperpolarizing pulse. Any cells with more than 15% change in either measure during control periods between drug response measurements were eliminated from the study. Agonists were applied via pressure ejection (Smart Squirt, Automate, Inc.), 60 s per application, followed by 30 s of control aCSF, from a 250 μm tip placed within 350 μm of the recorded cell. Agonist solutions were loaded into the Smart Squirt at concentrations at least 10x saturating so that even with diffusion a saturating concentration of agonist would reach the recorded cell.

Agonists, antagonists, ATP, GTP, and all other chemicals were obtained from Sigma or Tocris Bioscience. KOR antagonists were provided by BlackThorn Therapeutics, Inc.

### *In vitro* pharmacology at rat KOR

Cellular antagonist effects of BTRX-335140 and BTRX-395750 (0.3 nM – 0.3 μM) were assessed in a rat recombinant CHO cell line using a cAMP-based time-resolved FRET assay (Eurofins Cerep, France). Results were calculated as a percent inhibition following application of the KOR agonist, (-)-U50,488 (3 nM).

### Data analysis

Each agonist response was calculated as the difference in the *I*_holding_ between the 2 min just prior to the agonist application and the 30 s around the peak response. Results are presented as mean ± SEM. Violin plots were constructed from the kernel density estimate of the data, where the bandwidth of the kernel was set to the range of the data in the plot divided by 10. Over 90% of concentration response experiments and wash-in measurements were made blind to antagonist identity. Concentration response data were fitted with the Hill equation to estimate IC_50_ values in IGOR Pro (Wavemetrics). Dose response data were also fit to a 4-parameter log-logistic dose response model using the drc package in R to estimate variance. Paired t-Tests (vassarstats.net) were conducted to compare within cell baseline responses to MOR and DOR agonists with responses to these agonists in the presence of the KOR antagonists. Statistical significance was determined as *p* < 0.05. Data are available on OSF (DOI 10.17605/OSF.IO/AURZ7).

## Results

### Concentration responses for the KOR antagonists

Responses of VTA neurons to pressure ejection application of a super-saturating concentration of the KOR agonist U-69,593 were measured in acute horizontal brain slices from rats using whole cell electrophysiology in voltage clamp configuration. KOR activation under these conditions activates K+ channels in many neurons, which in a voltage clamp mode results in an outward (positive) current deflection (Fig 1a). Approximately half of VTA dopamine neurons are hyperpolarized by KOR activation [1], therefore each cell was tested for a U-69,593 response, and those that responded with an outward current were used to measure the efficacy of an antagonist to block the response to subsequent re-application of U-69,593. In control experiments of repeated U-69,593 testing without addition of antagonists, we found no evidence for desensitization of the U-69,593 response in this preparation: the second responses were 124 ± 7% the magnitude of the first responses (n = 9). For BTRX-335140, we measured an IC_50_ of 1.3 ± 0.9 nM (Fig 1b). Both 10 and 100 nM completely blocked the U-69,593 responses. This is quite similar to our measurements in a heterologous system expressing rat KORs, where BTRX- 335140 had an IC_50_ of 3.2 nM for blocking inhibition of adenylyl cylcase by (-)-U50,488 (3 nM). For a structurally related compound in the same series, BTRX-395750, we measured an IC_50_ of 4.6 ± 0.9 nM (Fig 1b) at KOR in VTA neurons, a greater potency than was measured in the heterologous system (IC_50_ = 48 nM).

**Fig 1.**
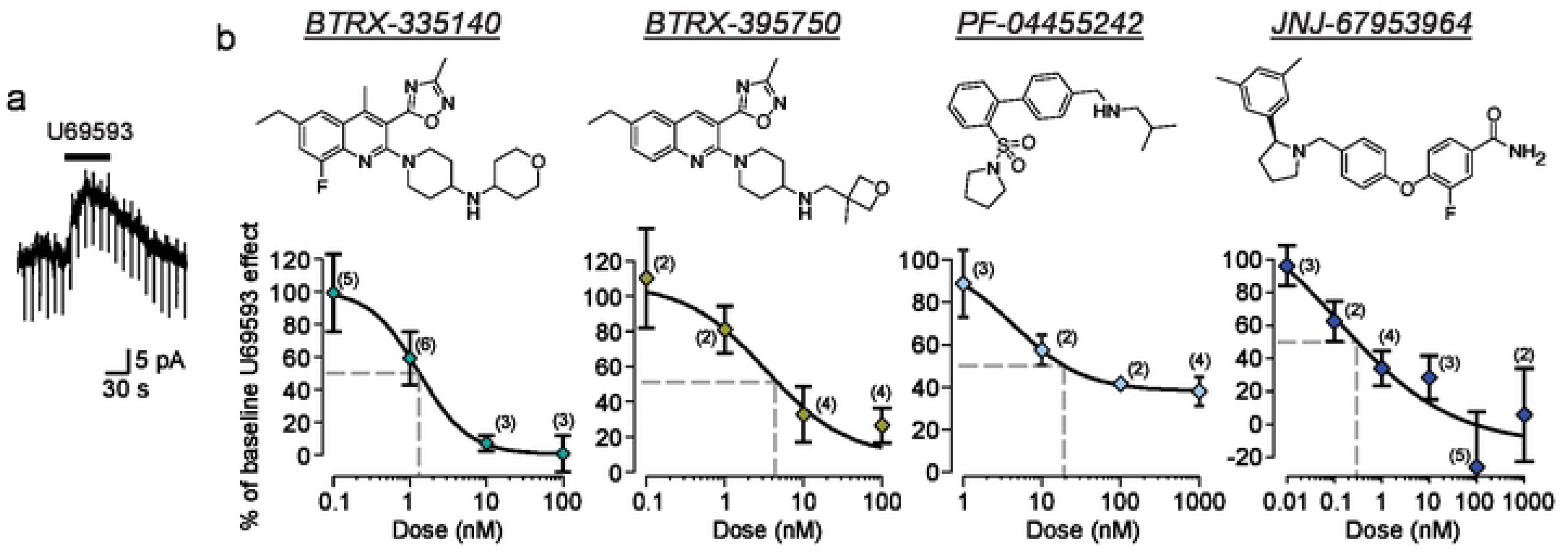
Concentration response relationships for blockade of U-69,593 induced K+ currents in VTA dopamine neurons by novel KOR antagonists. **a** Example voltage clamp recording of an outward current in a VTA neuron in response to pressure ejection of the KOR agonist U- 69,593. **b** Top row, structures of the 4 KOR antagonists tested. Bottom row, concentration response curves for blockade of U-69,593 responses for four recently developed KOR antagonists.

Although PF-04455242 is reported to be a full antagonist in heterologous systems [37], we found that it only partially blocked the U-69,593 responses in the electrophysiology assay (Fig 1b). We observed a maximal blockade of approximately 60% of the U-69,593 responses at both 100 nM and 1 μM PF-04455242. The calculated concentration that blocked 50% of the baseline U-69,593 response amplitude was 19.6 ± 16.0 nM. The concentration of PF-04455242 that produced half of the maximum effect for this antagonist is 4.3 nM. These data indicate that PF- 04455242 is a partial antagonist in this tissue.

We also studied the concentration response of JNJ-67953964, which yielded an IC_50_ of 0.3 ± 1.3 nM (Fig 1b). A surprising result in these experiments was that in the presence of 100 nM and 1 μM concentrations of JNJ-67953964, a subset of neurons responded to U-69,593 with inward currents instead of outward currents, indicating an off target effect of this compound.

We have previously determined that KOR activation specifically inhibits VTA dopamine neurons that project to the mPFC but not to the NAc [2]. To investigate the effects of KOR antagonism on this specific circuit, we measured the concentration response of BTRX-335140 blockade of U-69,593 responses in VTA dopamine neurons that project to the mPFC. The fluorescent tracer DiI was injected into the mPFC 6-8 days prior to *ex vivo* recordings, and was detected in somata prior to patching (Fig 2a). Consistent with prior observations [2], outward currents were observed in response to U-69,593 in VTA neurons that project to the mPFC, and these responses were blocked by BTRX-335140 (example in Fig 2a). In these selected neurons, the IC_50_ of BTRX-335140 was 2.2 ± 2.4 nM, similar to the IC_50_ determined among non-projection-selected neurons (Fig 2b).

**Fig 2.**
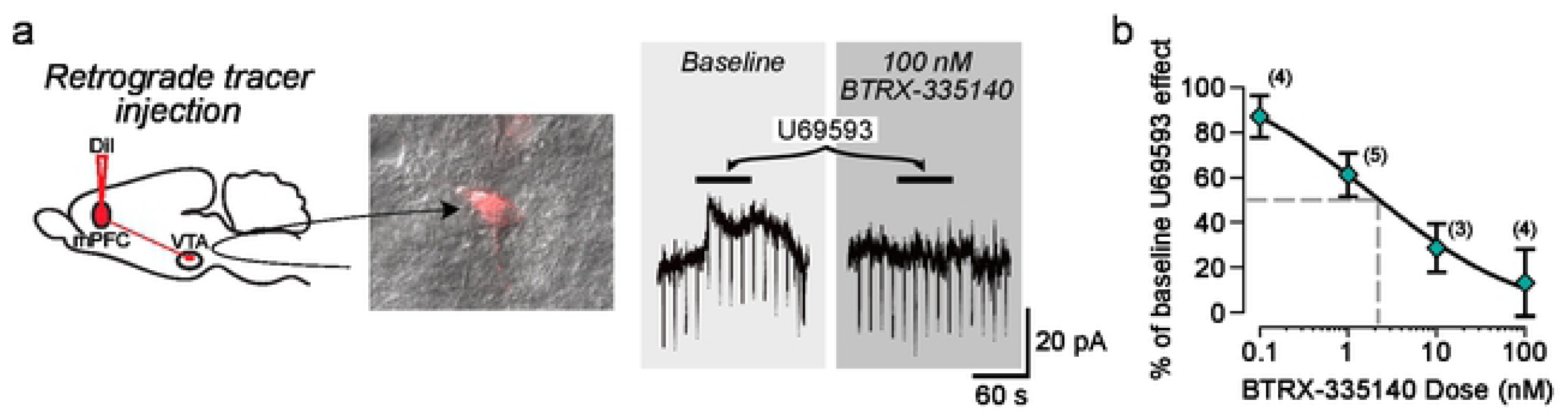
The KOR antagonist BTRX-335140 blocks U-69,593 responses in mPFC-projecting VTA dopamine neurons. **a** Example VTA neuron retrogradely labeled by stereotaxic injection of DiI into the mPFC (left). This neuron responded to U-69,593 application, and this outward current was completely blocked by the KOR antagonist BTRX-335140 (right). The concentration response relationship for BTRX-335140 specifically in VTA neurons that project to the mPFC (**b**) is similar to that measured among unselected neurons (Fig 1b).

### Selectivity of BTRX-335140 compared to JNJ-67953964 for KORs

To evaluate the selectivity of BTRX-335140 and JNJ-67953964 for KORs over MORs and DORs, we tested their ability to block selective agonist-induced responses at these two receptors in VTA neurons. Super-saturating concentrations of the MOR selective agonist DAMGO (10 μM) or the DOR selective agonist DPDPE (10 μM) were pressure ejected onto VTA neurons in the same manner as U-69,593. We have previously shown that DAMGO and DPDPE responses in VTA neurons in acute brain slices do not desensitize [3,4]. In responsive neurons, the agonist was re-applied after 10 nM of either KOR antagonist had been bath applied for at least 4 min. This concentration of BTRX-335140, which acted as a full antagonist to block the U-69,593 responses, did not affect the responses to DAMGO (n = 7, paired t-Test t = +1.05, df = 6, *p* = 0.33) or DPDPE (n = 5, paired t-Test t = +0.36, df = 4, *p* = 0.74; Fig 3a). By contrast, a 10 nM concentration of JNJ-67953964, which also completely blocked U-69,593 responses, consistently diminished responses to DAMGO (n = 5, paired t-Test t = +2.84, df = 4, *p* = 0.047) but not DPDPE (n = 3, paired t-Test t = +2.09, df = 2, *p* = 0.17; Fig 3b).

**Fig 3.**
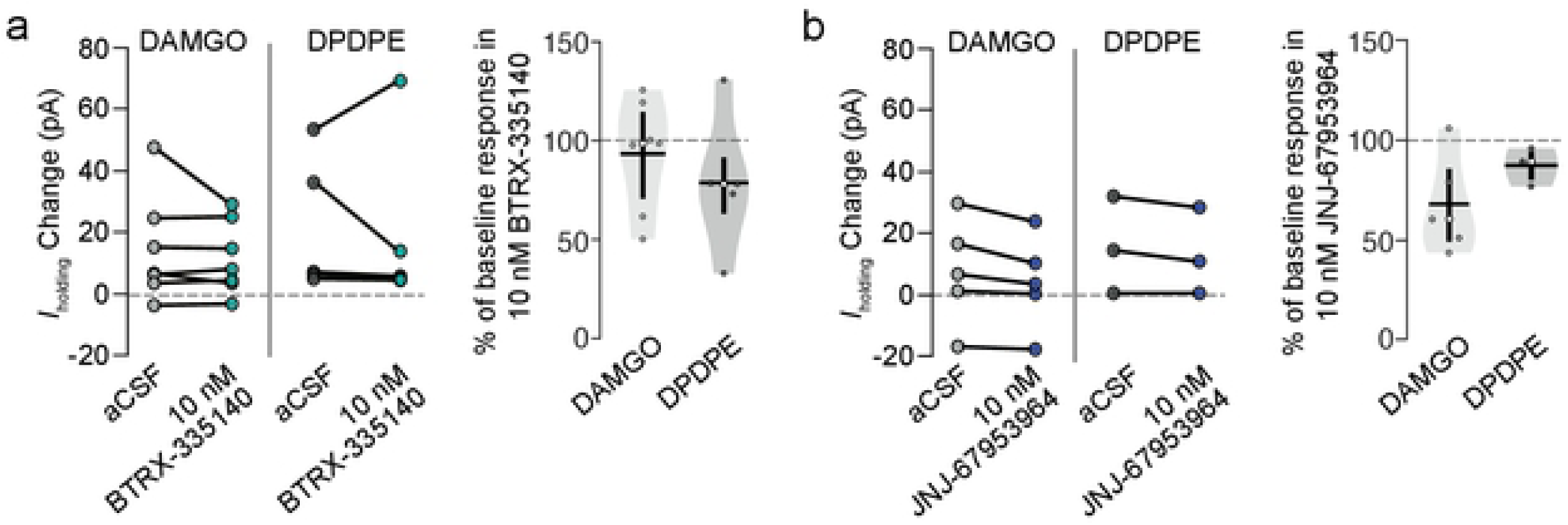
JNJ-67953964 is less selective for KOR over other opioid receptors compared to BTRX-335140 in VTA neurons. BTRX-335140 (**a**) and JNJ-67953964 (**b**) were tested for off target blockade of MOR or DOR antagonism with the agonists DAMGO and DPDPE, respectively. Neurons that responded with a change in holding current in response to DAMGO or DPDPE were re-tested with the agonist in the presence of 10 nM of the antagonist (right panel of **a** and **b**). The data were transformed showing the magnitude of the agonist response in the presence of the antagonist relative to the magnitude of the control, pre-antagonist response (left panel of **a** and **b**).

### Wash out of test antagonists

One major shortcoming of norBNI, the KOR antagonist used most broadly in preclinical studies, is the persistent effects of a single administration [22,26]. A short acting, selective antagonist is not only useful for clinical development, but also for experimental designs that require ligands to reverse rapidly enough for repeated administrations to have discernible effects. Here we measured whether responses to U-69,593 recovered after 10 or 20 min of washout of the novel antagonists. Concentrations of antagonists were chosen for their maximal blockade of U-69,593 effects in the concentration response experiments.

In each experiment, a baseline U-69,593 response was measured, then the antagonist was applied to the slice for at least 5 min. This interval and the antagonist concentration selected were conditions sufficient to completely block U-69,593 responses, as observed in concentration response experiments shown above. U-69,593 responses were then probed 10 and/or 20 minutes after antagonist washout commenced, depending on the duration of the stability the whole cell recording. As expected, application of a concentration of norBNI (100 nM) that we have previously used to either fully block [1] or reverse [38] U-69,593 effects in VTA neurons showed no reversal at 20 min of washout (Fig 4b). On the other hand, a 10 nM concentration of BTRX- 335140, sufficient to completely block U-69,593 actions (Figs 1b, 4a), showed complete washout within 10 min (Fig 4). Interestingly, BTRX-395750, which is structurally closely related to BTRX- 335140 (Fig 1b), did not show significant reversal with up to 20 min washout (Fig 4b). PF-04455242 also did not show substantial washout at 10 min, but did permit recovery of the U- 69,593 effect at 20 min (Fig 4b). JNJ-67953964 on average showed some apparent washout, but only to the extent that the inward currents produced by U-69,593 in the presence of JNJ- 67953964 in some neurons (Fig 1b) were not observed at either washout timepoint (Fig 4b).

**Fig 4.**
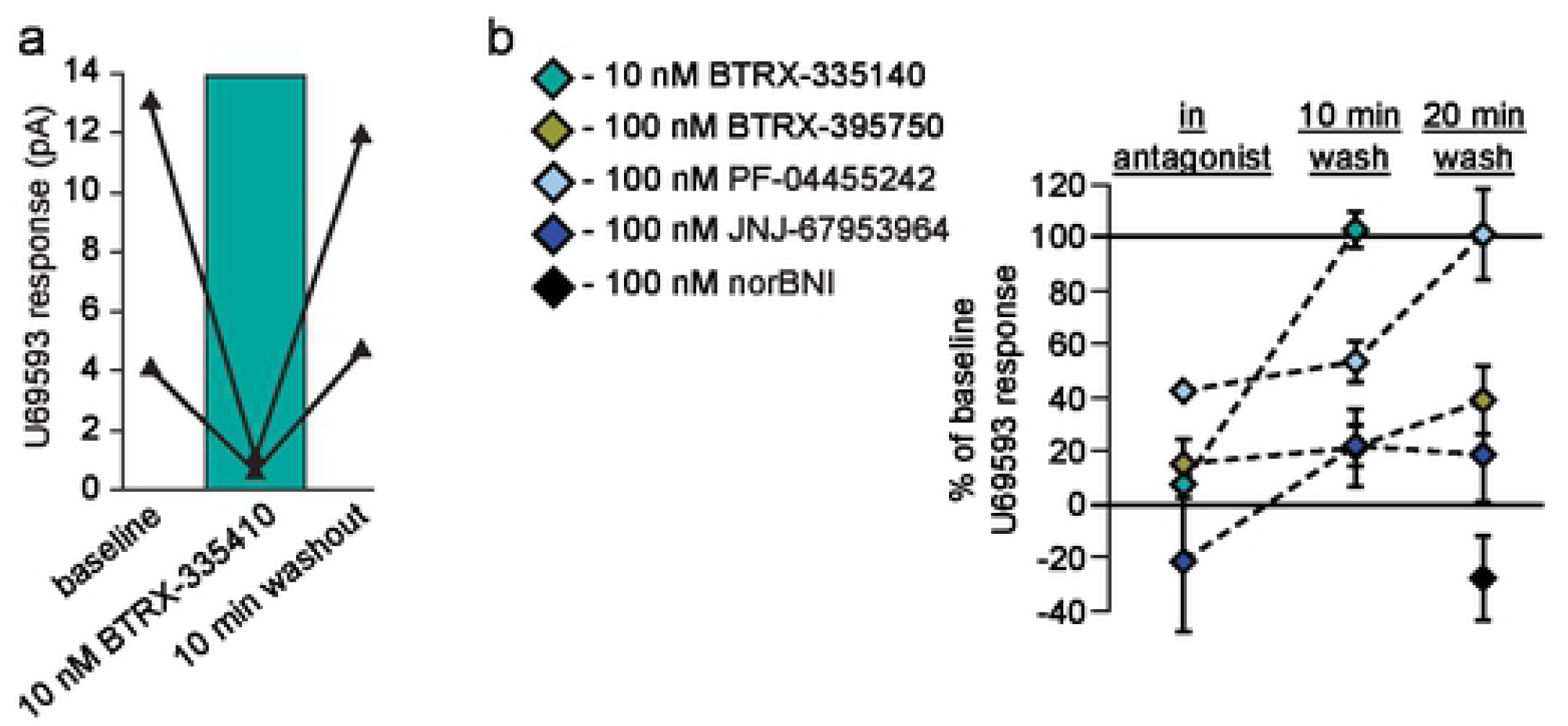
Washout of some novel KOR antagonists can be detected in VTA dopamine neurons. **a** Data from 2 different neurons showing that 10 nM BTRX-335140 completely blocked U-69,593 responses, and this blockade reversed following 10 min of antagonist washout. **b** Summary across antagonists showing that only BTRX-335140 reverses within 10 min. The partial antagonism of PF-04455242 reversed at 20 min.

### Wash in effects of test antagonists

In VTA slices, we have not observed effects of norBNI that would suggest either constitutive activity at the KOR or endogenous dynorphin release in midbrain tissue from naïve animals [1,38]. Therefore, neutral antagonists at the KOR would not be expected to drive any change in *I*_holding_ in voltage clamp recordings. Here we measured the change in *I*_holding_ following bath application of each antagonist at multiple concentrations. While BTRX-335140, BTRX- 395750, and JNJ-67953964 had no effect on baseline *I*_holding_, PF-04455242 did induce a shift in *I*_holding_ in a subset of neurons (Fig 5). Specifically, in a subset of VTA neurons, bath application of PF-04455242 caused an outward current at the 100 nM and 1 μM concentrations (Fig 5). This change in *I*_holding_ was accompanied by a decrease in input resistance, consistent with a channel opening (Fig 5a). Given the intracellular and extracellular solutions used here, and that the neurons were clamped at V_m_= −60 mV, this current is most likely K+ mediated, such as through a G protein coupled inwardly rectifying K+ channel (GIRK). GIRK is the typical postsynaptic coupled ion channel for opioid receptors, however, since every neuron tested in these wash in experiments responded to the KOR agonist U-69,593, yet only a subset of them show this response to PF-04455242, it is unlikely that these currents are due to activation of KORs.

**Fig 5.**
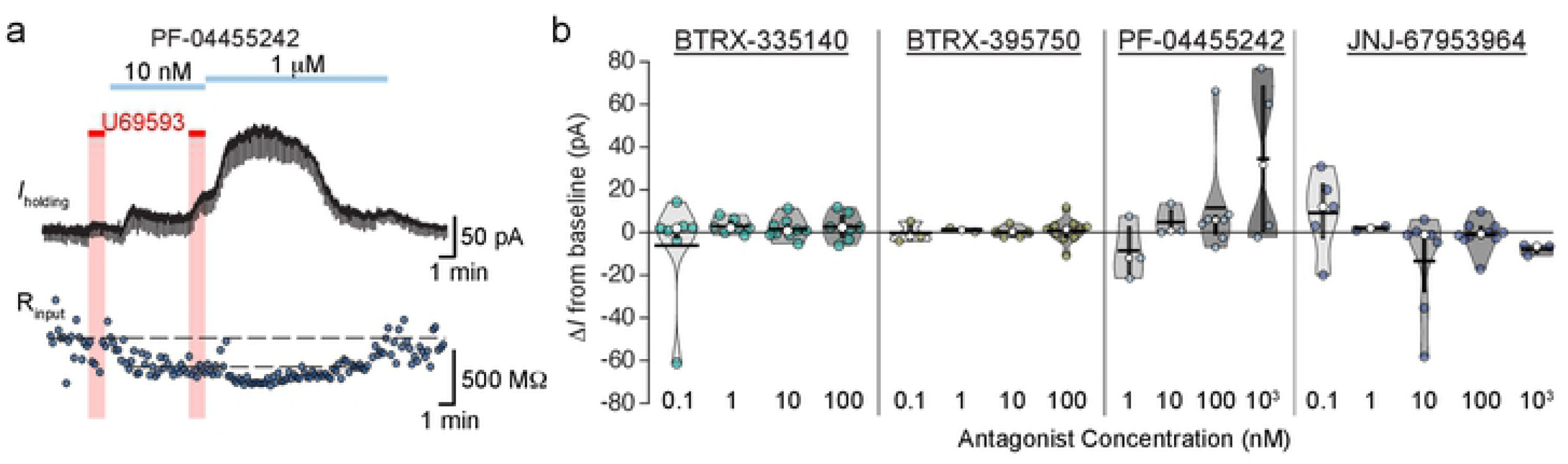
Among novel KOR antagonists, PF-04455242 stands out as activating a current. **a** Current (top) and input resistance (bottom) of example voltage clamp recording. This neuron responded to U-69,593 with a small outward current and decrease in input resistance. This neuron responded to PF-04455242 with a larger outward current and concurrent decrease in input resistance in a concentration dependent manner. **b** Summary of all changes in holding current in response to bath application of each KOR antagonist.

## Discussion

Here we tested the potency, selectivity, and reversibility of several recently developed KOR antagonists using electrophysiological measurements in acute rat brain slices. This preparation has the advantage of measuring ligand actions at receptors expressed endogenously in the mammalian brain. We selected the VTA dopaminergic neurons for profiling these molecules because this is a brain region where KOR agonists produce potent aversive actions, and because many other GPCRs are also expressed in VTA neurons, increasing the likelihood that off-target effects of these ligands could be detected. In fact, this approach did reveal some unpredicted properties of some of these putative KOR antagonists. Further, it enabled us to test for reversal during washout, neural modulation with wash in, and off target blockade of endogenously expressed MOR or DOR. BTRX-335140 in particular was potent (1.3 nM IC_50_), showed rapid reversal of KOR antagonism in washout, and lacked MOR or DOR antagonist effects at the concentrations tested. In preclinical studies [28], BTRX-335140 showed oral efficacy in target engagement measures and is currently in clinical development.

BTRX-395750 also exhibited <10 nM potency at the KOR in this preparation, however it showed no significant washout of KOR blockade up to 20 min. This is interesting given that this molecule is within the same chemical series as BTRX-335140 and they are structurally closely related. Understanding how the specific differences in the structures affect the molecules’ orientations in the KOR binding pocket may inform what receptor-ligand interactions contribute to receptor residence times for the KOR. This is particularly interesting given the anomalously long duration of action of not only norBNI, but also the nonmorphinan KOR antagonist JDTic [23].

PF-04455242 showed some unexpected results in the characterization studies performed here compared to previously described pharmacological properties [29,37]. First, we found it to only have partial antagonist action, with maximal blockade of the U-69,593 response plateauing at approximately 60%. We also found that PF-04455242 appears to activate a K+ current on a subset of the recorded cells. It is unlikely that these responses were due to activation of KOR, since all of the neurons tested with the compound initially responded to U-69,593. Consistent with these responses not being due to KOR activation, in HEK293 cells expressing KOR, PF- 04455242 did not generate *extracellular signal-regulated kinase* (ERK) phosphorylation and minimal *c-Jun N terminal kinase* (JNK) phosphorylation compared to other KOR antagonists [23]. This compound was reported to have only moderate binding selectivity for KOR over MOR in radioligand displacement studies in rat brain tissue [37]. Together, these observations indicate PF-04455242 is quite different from a neutral KOR selective antagonist.

JNJ-67953964 effects have been explored in a variety of preclinical and human studies. It is brain-penetrant and well tolerated in human studies [39–41], including in people in early abstinence from cocaine dependence [42]. In heterologous systems this compound acts as a neutral antagonist at KOR and at higher concentrations also blocks MOR [25]. However in our acute brain slice preparation, we detected an unexpected effect at the KOR in a subset of neurons, wherein JNJ-67953964 at concentrations ≥ 100 nM switched the expected outward current driven by KOR activation to an inward current. This change in signaling appeared to wash out acutely within 10 min. It is unclear if this change in signaling is due to a JNJ-67953964 interaction with KOR, or through an action at another receptor. Since this switch was not uniform across all neurons tested, a direct action at KOR seems insufficient to explain how this effect would only be observed in a subset of neurons. While this effect is unique compared to the other antagonists investigated here, since it was only observed at relatively high concentrations, it may not be a major concern following systemic drug administration. On the other hand, it may be a confound in animal studies where JNJ-67953964 is centrally administered e.g. [43]. In our selectivity experiments, we also saw modest blockade of our DAMGO induced effects at just 10 nM JNJ-67953964. While some MOR antagonism has been reported for this compound previously [25,40], we were surprised to detect it at this concentration, which did not achieve full KOR blockade in this preparation. Together, these observations indicate potentially important, unanticipated properties of JNJ-67953964.

Here we characterized the pharmacological properties of putative reversible, KOR selective antagonists using whole cell recordings in acute brain slices. Our results provide novel and in some cases surprising findings on how these compounds work in individual neurons that endogenously express KORs. In particular, this approach enabled us to (1) better identify the activity of these compounds at the KOR and (2) identify off-target effects of these compounds that were not previously understood. Not only does this work reveal the functional differences between these compounds that inform their use in both preclinical studies and clinical development, this study provides direct evidence that electrophysiology in the acute brain slice preparation enables detection of molecular properties of novel compounds that are not easily detected in conventional heterologous receptor expression systems used for drug screening.

## Acknowledgements

We thank Catriona Miller and Kasra Mansourian for technical support and Howard Fields for helpful comments on the manuscript.

## References

1. Margolis EB, Hjelmstad GO, Bonci A, Fields HL. Kappa-opioid agonists directly inhibit midbrain dopaminergic neurons. J Neurosci. 2003;23:9981–9986.

2. Margolis EB, Lock H, Chefer VI, Shippenberg TS, Hjelmstad GO, Fields HL. Kappa opioids selectively control dopaminergic neurons projecting to the prefrontal cortex. Proc Natl Acad Sci U A. 2006;103:2938–2942.

3. Margolis EB, Hjelmstad GO, Fujita W, Fields HL. Direct bidirectional μ-opioid control of midbrain dopamine neurons. J Neurosci. 2014;34:14707–14716.

4. Margolis EB, Fujita W, Devi LA, Fields HL. Two delta opioid receptor subtypes are functional in single ventral tegmental area neurons, and can interact with the mu opioid receptor. Neuropharmacology. 2017;123:420–432.

5. Margolis EB, Mitchell JM, Hjelmstad GO, Fields HL. A novel opioid receptor-mediated enhancement of GABAA receptor function induced by stress in ventral tegmental area neurons. J Physiol. 2011;589:4229–4242.

6. Kallupi M, Wee S, Edwards S, Whitfield TW, Oleata CS, Luu G, et al. Kappa opioid receptor-mediated dysregulation of gamma-aminobutyric acidergic transmission in the central amygdala in cocaine addiction. Biol Psychiatry. 2013;74:520–528.

7. Bajo M, Madamba SG, Roberto M, Siggins GR. Acute morphine alters GABAergic transmission in the central amygdala during naloxone-precipitated morphine withdrawal: role of cyclic AMP. Front Integr Neurosci. 2014;8:45.

8. McPherson KB, Leff ER, Li M-H, Meurice C, Tai S, Traynor JR, et al. Regulators of G-Protein Signaling (RGS) Proteins Promote Receptor Coupling to G-Protein-Coupled Inwardly Rectifying Potassium (GIRK) Channels. J Neurosci Off J Soc Neurosci. 2018;38:8737–8744.

9. Walsh SL, Strain EC, Abreu ME, Bigelow GE. Enadoline, a selective kappa opioid agonist: comparison with butorphanol and hydromorphone in humans. Psychopharmacology (Berl). 2001;157:151–162.

10. Pfeiffer A, Brantl V, Herz A, Emrich HM. Psychotomimesis mediated by kappa opiate receptors. Science. 1986;233:774–776.

11. Tejeda HA, Bonci A. Dynorphin/kappa-opioid receptor control of dopamine dynamics: Implications for negative affective states and psychiatric disorders. Brain Res. 2018. 19 September 2018. https://doi.org/10.1016/j.brainres.2018.09.023.

12. Land BB, Bruchas MR, Lemos JC, Xu M, Melief EJ, Chavkin C. The dysphoric component of stress is encoded by activation of the dynorphin kappa-opioid system. J Neurosci. 2008;28:407–414.

13. Bruchas MR, Land BB, Aita M, Xu M, Barot SK, Li S, et al. Stress-induced p38 mitogen-activated protein kinase activation mediates kappa-opioid-dependent dysphoria. J Neurosci. 2007;27:11614–11623.

14. McLaughlin JP, Marton-Popovici M, Chavkin C. Kappa opioid receptor antagonism and prodynorphin gene disruption block stress-induced behavioral responses. J Neurosci. 2003;23:5674–5683.

15. Walker BM, Zorrilla EP, Koob GF. Systemic κ-opioid receptor antagonism by nor-binaltorphimine reduces dependence-induced excessive alcohol self-administration in rats. Addict Biol. 2011;16:116–119.

16. Chavkin C, Koob GF. Dynorphin, Dysphoria, and Dependence: the Stress of Addiction. Neuropsychopharmacology. 2016;41:373–374.

17. Chartoff E, Sawyer A, Rachlin A, Potter D, Pliakas A, Carlezon WA. Blockade of kappa opioid receptors attenuates the development of depressive-like behaviors induced by cocaine withdrawal in rats. Neuropharmacology. 2012;62:167–176.

18. Navratilova E, Ji G, Phelps C, Qu C, Hein M, Yakhnitsa V, et al. Kappa opioid signaling in the central nucleus of the amygdala promotes disinhibition and aversiveness of chronic neuropathic pain. Pain. 2019;160:824–832.

19. Wells AM, Ridener E, Bourbonais CA, Kim W, Pantazopoulos H, Carroll FI, et al. Effects of Chronic Social Defeat Stress on Sleep and Circadian Rhythms Are Mitigated by Kappa-Opioid Receptor Antagonism. J Neurosci Off J Soc Neurosci. 2017;37:7656–7668.

20. Bruchas MR, Yang T, Schreiber S, Defino M, Kwan SC, Li S, et al. Long-acting kappa opioid antagonists disrupt receptor signaling and produce noncompetitive effects by activating c-Jun N-terminal kinase. J Biol Chem. 2007;282:29803–29811.

21. Carroll FI, Carlezon WA. Development of κ opioid receptor antagonists. J Med Chem. 2013;56:2178–2195.

22. Patkar KA, Wu J, Ganno ML, Singh HD, Ross NC, Rasakham K, et al. Physical presence of nor-binaltorphimine in mouse brain over 21 days after a single administration corresponds to its long-lasting antagonistic effect on κ-opioid receptors. J Pharmacol Exp Ther. 2013;346:545–554.

23. Melief EJ, Miyatake M, Carroll FI, Béguin C, Carlezon WA, Cohen BM, et al. Duration of action of a broad range of selective κ-opioid receptor antagonists is positively correlated with c-Jun N-terminal kinase-1 activation. Mol Pharmacol. 2011;80:920–929.

24. Schattauer SS, Bedini A, Summers F, Reilly-Treat A, Andrews MM, Land BB, et al. Reactive oxygen species (ROS) generation is stimulated by κ opioid receptor activation through phosphorylated c-Jun N-terminal kinase and inhibited by p38 mitogen-activated protein kinase (MAPK) activation. J Biol Chem. 2019;294:16884–16896.

25. Rorick-Kehn LM, Witkin JM, Statnick MA, Eberle EL, McKinzie JH, Kahl SD, et al. LY2456302 is a novel, potent, orally-bioavailable small molecule kappa-selective antagonist with activity in animal models predictive of efficacy in mood and addictive disorders. Neuropharmacology. 2014;77:131–144.

26. Munro TA, Berry LM, Van’t Veer A, Béguin C, Carroll FI, Zhao Z, et al. Long-acting κ opioid antagonists nor-BNI, GNTI and JDTic: pharmacokinetics in mice and lipophilicity. BMC Pharmacol. 2012;12:5.

27. Urbano M, Guerrero M, Rosen H, Roberts E. Antagonists of the kappa opioid receptor. Bioorg Med Chem Lett. 2014;24:2021–2032.

28. Guerrero M, Urbano M, Kim E-K, Gamo AM, Riley S, Abgaryan L, et al. Design and Synthesis of a Novel and Selective Kappa Opioid Receptor (KOR) Antagonist (BTRX-335140). J Med Chem. 2019;62:1761–1780.

29. Verhoest PR, Basak AS, Parikh V, Hayward M, Kauffman GW, Paradis V, et al. Design and discovery of a selective small molecule κ opioid antagonist (2-methyl-N-((2’-(pyrrolidin-1-ylsulfonyl)biphenyl-4-yl)methyl)propan-1-amine, PF-4455242). J Med Chem. 2011;54:5868–5877.

30. Margolis EB, Karkhanis AN. Dopaminergic cellular and circuit contributions to kappa opioid receptor mediated aversion. Neurochem Int. 2019;129:104504.

31. Fallon JH, Leslie FM, Cone RI. Dynorphin-containing pathways in the substantia nigra and ventral tegmentum: a double labeling study using combined immunofluorescence and retrograde tracing. Neuropeptides. 1985;5:457–460.

32. Chou TC, Lee CE, Lu J, Elmquist JK, Hara J, Willie JT, et al. Orexin (Hypocretin) Neurons Contain Dynorphin. J Neurosci. 2001;21:RC168–RC168.

33. Fadel J, Deutch AY. Anatomical substrates of orexin–dopamine interactions: lateral hypothalamic projections to the ventral tegmental area. Neuroscience. 2002;111:379–387.

34. Knoll AT, Carlezon WA. Dynorphin, stress, and depression. Brain Res. 2010;1314:56–73.

35. Koob GF. The dark side of emotion: the addiction perspective. Eur J Pharmacol. 2015;753:73–87.

36. Ehrich JM, Messinger DI, Knakal CR, Kuhar JR, Schattauer SS, Bruchas MR, et al. Kappa Opioid Receptor-Induced Aversion Requires p38 MAPK Activation in VTA Dopamine Neurons. J Neurosci. 2015;35:12917–12931.

37. Grimwood S, Lu Y, Schmidt AW, Vanase-Frawley MA, Sawant-Basak A, Miller E, et al. Pharmacological characterization of 2-methyl-N-((2’-(pyrrolidin-1-ylsulfonyl)biphenyl-4-yl)methyl)propan-1-amine (PF-04455242), a high-affinity antagonist selective for κ-opioid receptors. J Pharmacol Exp Ther. 2011;339:555–566.

38. Margolis EB, Hjelmstad GO, Bonci A, Fields HL. Both kappa and mu opioid agonists inhibit glutamatergic input to ventral tegmental area neurons. J Neurophysiol. 2005;93:3086–3093.

39. Lowe SL, Wong CJ, Witcher J, Gonzales CR, Dickinson GL, Bell RL, et al. Safety, tolerability, and pharmacokinetic evaluation of single- and multiple-ascending doses of a novel kappa opioid receptor antagonist LY2456302 and drug interaction with ethanol in healthy subjects. J Clin Pharmacol. 2014;54:968–978.

40. Rorick-Kehn LM, Witcher JW, Lowe SL, Gonzales CR, Weller MA, Bell RL, et al. Determining pharmacological selectivity of the kappa opioid receptor antagonist LY2456302 using pupillometry as a translational biomarker in rat and human. Int J Neuropsychopharmacol. 2014;18.

41. Naganawa M, Dickinson GL, Zheng M-Q, Henry S, Vandenhende F, Witcher J, et al. Receptor Occupancy of the κ-Opioid Antagonist LY2456302 Measured with Positron Emission Tomography and the Novel Radiotracer 11C-LY2795050. J Pharmacol Exp Ther. 2016;356:260–266.

42. Reed B, Butelman ER, Fry RS, Kimani R, Kreek MJ. Repeated Administration of Opra Kappa (LY2456302), a Novel, Short-Acting, Selective KOP-r Antagonist, in Persons with and without Cocaine Dependence. Neuropsychopharmacol Off Publ Am Coll Neuropsychopharmacol. 2018;43:739–750.

43. Custodio-Patsey L, Donahue RR, Fu W, Lambert J, Smith BN, Taylor BK. Sex differences in kappa opioid receptor inhibition of latent postoperative pain sensitization in dorsal horn. Neuropharmacology. 2019:107726.

